# Processes driving nocturnal transpiration and implications for estimating land evapotranspiration

**DOI:** 10.1101/017202

**Authors:** Vίctor Resco de Dios, Jacques Roy, Juan Pedro Ferrio, Josu G. Alday, Damien Landais, Alexandru Milcu, Arthur Gessler

**Affiliations:** Department of Crop and Forest Sciences-AGROTECNIO Center, Universitat de Lleida, 25198 Lleida, Spain.; Hawkesbury Institute for the Environment, University of Western Sydney, Richmond, NSW 2753, Australia.; Ecotron Européen de Montpellier, UPS 3248, CNRS, Campus Baillarguet, 34980, Montferrier-sur-Lez, France.; School of Environmental Sciences, University of Liverpool, Liverpool, L69 3GP, UK.; CNRS, Centre d’Ecologie Fonctionnelle et Evolutive (CEFE, UMR-5175), 1919, route de Mende, 34293, Montpellier, France.; Swiss Federal Institute for Forest, Snow and Landscape Research WSL Long-term Forest Ecosystem Research (LWF), 8903 Birmensdorf, Switzerland.; Institute for Landscape Biogeochemistry, Leibniz-Centre for Agricultural Landscape Research (ZALF), 15374 Müncheberg, Germany.

## Abstract

Evapotranspiration is a major component of the water cycle, yet only daytime transpiration is currently considered in Earth system and agricultural sciences. This contrasts with physiological studies where 25% or more of water losses have been reported to occur occurring overnight at leaf and plant scales. This gap probably arose from limitations in techniques to measure nocturnal water fluxes at ecosystem scales, a gap we bridge here by using lysimeters under controlled environmental conditions. The magnitude of the nocturnal water losses (12-23% of daytime water losses) in row-crop monocultures of bean (annual herb) and cotton (woody shrub) would be globally an order of magnitude higher than documented responses of global evapotranspiration to climate change (51-98 *vs.* 7-8 mm yr^−1^). Contrary to daytime responses and to conventional wisdom, nocturnal transpiration was not affected by previous radiation loads or carbon uptake, and showed a temporal pattern independent of vapour pressure deficit or temperature, because of endogenous controls on stomatal conductance via circadian regulation. Our results have important implications from large-scale ecosystem modelling to crop production: homeostatic water losses justify simple empirical predictive functions, and circadian controls show a fine-tune control that minimizes water loss while potentially increasing posterior carbon uptake.

---

Global evapotranspiration is estimated to return annually 60% of total precipitation to the atmosphere^1^. Daytime transpiration dominates in current estimates of global evapotranspiration^2^, but nocturnal transpiration is largely unaccounted for^3^. At the leaf level, stomata have traditionally been assumed to close during the night and, in combination with low evaporative demand, were thought to cause a negligible water vapour flux^3^. However, there is growing evidence at the leaf and plant scales that incomplete stomatal closure and subsequent transpiration overnight are widespread and significant^4^. Recent studies estimate that the equivalent to 10-15% of daytime leaf and plant water losses occur overnight^4^, with values reaching 25-30% in desert and savanna plants^5,6^, and with some extreme species reaching higher night-time than daytime transpiration^7^.

Much of the current discussion on global evapotranspiration is focused on changes in the intensity of the water cycle under global warming and its impacts on crop productivity^8,9^. If the equivalent to 10-15% of daytime transpiration is lost overnight also at ecosystem scales, nocturnal transpiration could have a strong impact on global evapotranspiration, much higher than the impacts of documented and predicted accelerations or decelerations of the global water cycle (typically ∼1-2% of global evapotranspiration)^8,9^. However, quantifying and understanding the drivers of nocturnal water loss in ecosystems is challenging because of limitations in current techniques. In addition to the difficulties in partitioning evaporation from transpiration^2^: i) negative net radiation and condensation are not well represented in evapotranspiration models^10^; ii) direct measurements of latent heat flux overnight with flux towers are problematic due to low atmospheric turbulence^11^; iii) remote sensing estimates from reflectance cannot be obtained^12^; and iv) up-scaling measurements of sap flux is limited to woody species and complicated by water refilling in stem capacitors after daytime water losses^13-15^.

Here we used 2 m^2^ lysimeters inside climate controlled macrocosms^16^ to obtain high-accuracy estimates of nocturnal transpiration over entire ecosystems (a dark plastic cover prevented evaporation from the soil), using row-crop monocultures of bean (*Phaseolus vulgaris*), an annual herb; and cotton (*Gossypium hirsutum*), a perennial shrub, as our model ecosystems. Our ultimate goals were to test whether nocturnal transpiration would be a significant component of whole ecosystem transpiration, and to examine the underlying mechanisms to provide guidance on how to address this process in large-scale water cycle studies and in agricultural development. We chose to focus on three important mechanisms that have been hypothesized to drive nocturnal transpiration, but only seldom explored:

1. We tested the influence of previous day radiation and daily carbon uptake on the magnitude of nocturnal transpiration. It has been hypothesized that nocturnal transpiration is influenced by a carry-over of daytime processes and, in particular, that photosynthesis regulates conductance in the following night by influencing carbohydrate supply, the necessary osmoticant for stomatal regulation^17,18^. We thus quantified nocturnal conductance and water losses after exposing the crops to different radiation environments, representative of a summer sunny day and of an autumn cloudy day, and expected to observe larger nocturnal water losses under high irradiance and carbon uptake.
2. We sought to determine whether the temporal pattern of nocturnal transpiration was driven by direct physiological responses to changes in vapour pressure deficit and temperature and/or by endogenous stomatal regulation. A major potential hindrance for modelling nocturnal transpiration is that different day-*vs*-night stomatal behaviours have been suggested, which would imply that applying models based on daytime parameterizations to nocturnal time periods would be problematic^5^. For instance, night-time stomatal conductance either does not respond to, or responds positively to, vapour pressure deficit in some species, contrary to the daytime trend^4,19^. Moreover, stomata tend to show higher conductance later than earlier in the night for a given level of vapour pressure deficit^20^ which would indicate that, in addition to direct responses to environmental variation, endogenous processes also affect nocturnal stomatal conductance. We thus expected to find weak direct responses to vapour pressure deficit variation^21^ and an interaction between direct physiological responses and endogenous controls^22^.
3. We tested the hypothesis that circadian regulation was the mechanism underlying endogenous patterns of stomatal conductance. Endogenous regulation has often been attributed to the circadian clock^4,19^ but, to show circadian regulation, one would need to experimentally maintain constant environmental conditions under prolonged (>24 h) darkness. As far as we are aware, this has only been done by one study^23^ so far and under CO_2_-free air, which potentially affected stomatal behaviour.

## Results

### Significant nocturnal transpiration unaltered by photosynthesis and radiation

We observed significant and substantial losses of water overnight. When photosynthetically active radiation, temperature and vapour pressure deficit mimicked the average pattern of an August sunny day in Montpellier (0/1,500 μmol m^−2^ s^−1^, min/max, 19/28 °C and 0.4/1.7 kPa, with 9 and 15 hours of night-time and of daytime, respectively), bean and cotton crops lost 0.8 and 0.6 mm d^−1^ during the night, which corresponded to 12% of daytime transpiration (Fig. 1).

**Figure 1:**
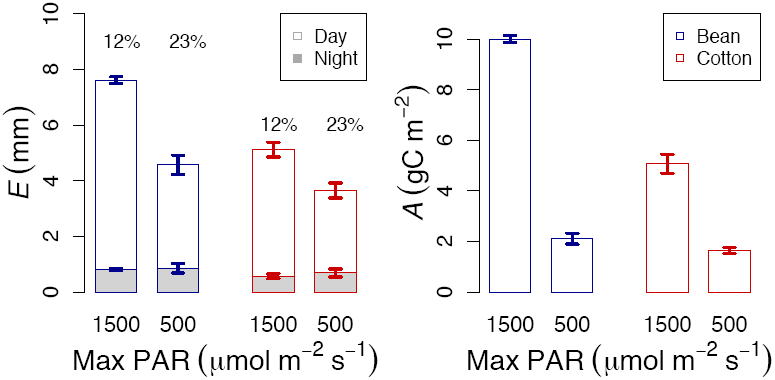
Transpiration and Carbon assimilation under high and low radiation. Under high radiation (maximum Photosynthetically Active Radiation, PAR, of 1,500 μmol m^−2^ s^−1^, and 15/9 h of daytime/night-time) transpiration (*E*) and Carbon assimilation (*A*) were higher than under reduced radiation (maximum PAR of 500 μmol m^−2^ s^−1^, and 12/12 h of daytime/night-time). However, the amount of nocturnal transpiration remained constant with radiation loads, indicating no impact of radiation and photosynthesis. The differing responses of daytime and night-time *E* to radiation led to an increase in the contribution of nocturnal *E*, relative to daytime *E*, from 12% to 23% as radiation loads decreased. Each bar indicates mean values of 6 macrocosms per species over 3-4 days and errors are SE.

There was no significant change in nocturnal transpiration when recreating the radiation load of an autumn cloudy day. We reduced the radiation loads by jointly increasing the duration of the dark period (12h of night-time and of daytime) and by allowing a maximum PAR of only 500 μmol m^−2^ s^−1^. Although the amount of nocturnal transpiration was slightly higher under reduced radiation loads, differences were significant neither in bean (linear mixed effects analyses, *P* = 0.50, *F* = 0.47) nor in cotton (linear mixed effects analyses, *P* = 0.47, *F* = 0.54; Fig. 1). We tested if this was an artefact arising from the different durations of the dark periods by comparing water losses during the 9 h of darkness that occurred when PAR varied between 0/1,500 μmol m^−2^ s^−1^ (21.00 – 06.00h) with those after 9 hours of darkness under 0/500 μmol m^−2^ s^−1^ (also 21.00 – 06.00h, that is, excluding the first 3 h of darkness in the simulated autumn cloudy day, and including only the last 9 h). Here, transpiration under reduced radiation was slightly lower than under high radiation, but differences were not significant (linear mixed effects analyses, *P* = 0.16, *F* = 2.98 in bean and *P* = 0.95, *F* = 0.005), indicating that the lack of differences across radiation environments was not affected by the duration of the dark period. Under low (autumn-like) radiation, the equivalent to 23% of daytime water losses in bean and cotton occurred during the night, due to significant declines in daytime water loss (Fig. 1).

Canopy conductance was also unaffected by radiation loads. We derived a proxy for canopy conductance (from the ratio between transpiration and vapour pressure deficit) and observed no differences across radiation environments (linear mixed effects analyses, *P* = 0.18, *F* = 1.83 in bean and *P* = 0.82, *F* = 0.05 in cotton). However, there were marked differences in integrated carbon assimilation across radiation environments (Fig. 1).

### Constancy in nocturnal transpiration co-driven by exogenous and endogenous processes

Rates of nocturnal transpiration remained constant, despite a 62% change in vapour pressure deficit between dusk and dawn (1.2 – 0.45 kPa, Fig. 2). The largest decline in vapour pressure deficit occurred during the first hours of the night (until 24.00 h) and was then followed by a period with limited variation in vapour pressure deficit and an increase in the proxy for canopy conductance (Fig. 2). The interaction between the initial decline in vapour pressure deficit and the posterior increase in canopy conductance led to no significant changes in the temporal pattern of nocturnal canopy transpiration (Fig. 2).

**Figure 2:**
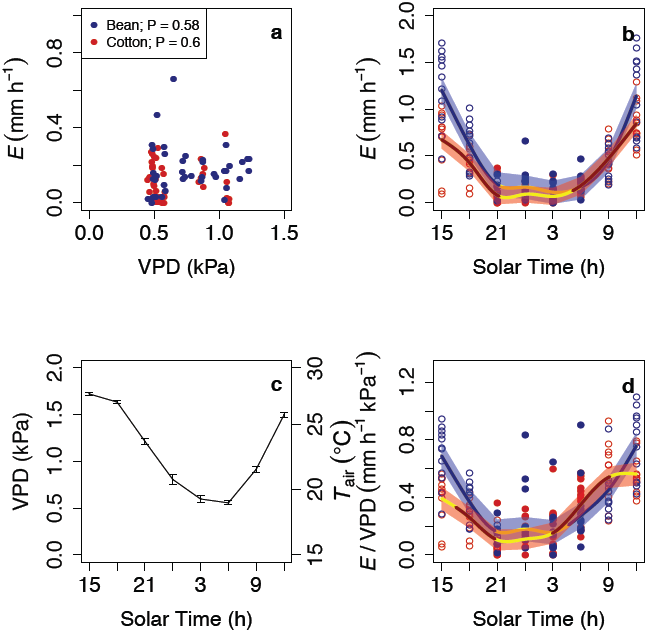
Temporal patterns of transpiration and drivers. Transpiration (*E*) remained constant overnight (**b**), and was not driven by vapour pressure deficit (VPD, **a**). The interaction between a declining VPD and air temperature (*T*_air_) early in the night (**c**) and an increase in canopy conductance (as indicated by *E*/VPD) later in the night (**d**) led to non-significant variation in *E* (**a**). Empty and filled dots mean daytime and night-time values, respectively. Lines (and shaded error intervals) indicate the prediction (and SE) of Generalized Additive Mixed Model (GAMM) fitting separately for each species (some lines may overlap), and significant temporal variation occurs only when the GAMM best-fit line is not yellow. Each dot represents the mean across 3-4 measuring nights for each of the 6 macrocosms per species and for each of the two radiation treatments. No differences in VPD across experiments and species existed (at *P* < 0.05), and only the mean pattern (and SE) is shown. A single line is fitted for both radiation environments given lack of differences in night-time rates (see text).

### Circadian regulation of endogenous canopy conductance

The late night increase in canopy conductance was independent of environmental variation (Fig. 2) and driven solely by endogenous circadian regulation of stomatal conductance (Fig. 3). Variation in nocturnal vapour pressure deficit after 03.00h was minimal, while canopy conductance significantly increased in both species overnight (Fig. 2), indicating significant endogenous stomatal regulation. We further observed that, at the leaf level, the late night increase in stomatal conductance was maintained even when we experimentally held constant levels of temperature (19°C), vapour pressure deficit (0.46 kPa) and light (0 μmol m^−2^ s^−1^) for 30 hours (Fig. 3). Stomatal conductance under prolonged darkness showed a cycle with a ∼24 h period (Fig. 3), and can thus be only attributed to circadian regulation.

**Fig. 3:**
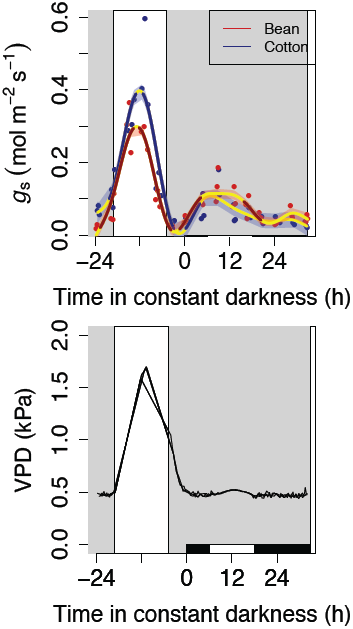
Circadian regulation of stomatal conductance in the dark. Environmental conditions of temperature and vapour pressure deficit (VPD) mimicked an average August day in Montpellier, with PAR recreating an autumn cloudy day (first 24 h shown), and remained constant for the following 30 h starting at solar midnight. The grey (white) background indicates when PAR was at (above) 0 μmol m^−2^ s^−1^. The white and black rectangles at the base indicate the subjective day and subjective night, respectively, under constant conditions. Points represent average values for each of three replicate macrocosms, and lines (and shaded error intervals) indicate the prediction (and SE) of Generalized Additive Mixed Model (GAMM) fitting separately for each species (some lines may overlap). Significant temporal variation (GAMM best-fit line portions not yellow) under constant conditions, with a ∼24 h cycle, can be fully attributed to circadian action. Lack of variation after 24h is likely due to carbon starvation. No differences in VPD across experiments and species existed (at *P* < 0.05) and only mean patterns are shown.

## Discussion

If our observations that nocturnal ecosystem transpiration account for the equivalent to 12-23% of daytime transpiration can be generalized (which is consistent with the wealth of studies at leaf and plant levels^4^), and accepting that 45%^2^ of the 950 mm^24^ of annual precipitation on Earth are transpired during the daytime, nocturnal transpiration would additionally return 51-98 mm yr^−1^ globally to the atmosphere. Global evapotranspiration has been reported to increase by 7 mm yr^−1^ between 1982-1997, and to decrease by 8 mm yr^−1^ during 1998-2008, presumably because of global warming-induced soil moisture shortages^6^. Nocturnal transpiration would therefore have an impact on global evapotranspiration an order of magnitude higher than current changes in the intensity of the evapotranspiration cycle resulting from global warming.

Under our conditions of optimal watering, the same amount of water was lost overnight under high and low radiation. This indicates that neither photosynthesis nor carbohydrate availability influence the amount of water lost overnight. However, when experimentally keeping continuous darkness for 30h, we observed a very marked cycle in stomatal conductance only during the first 24 hours, with constant and low values afterwards (Fig. 3), presumably because of carbon starvation after >24 hours of constant darkness. Our results thus help refine previous hypotheses^18^, indicating that photosynthesis and carbohydrates may impact nocturnal water fluxes in the field only under conditions that lead to carbon starvation (such as an extreme drought^25^). This implies that we can expect constancy in nocturnal transpiration regardless of daytime conditions, for as long as no important environmental stress occurs. If confirmed by additional studies, this should simplify large-scale Earth System modelling efforts as nocturnal transpiration could then be included empirically as a fixed amount, obtained from a fixed fraction of diurnal transpiration in sunny days.

Another step necessary for understanding how to represent nocturnal transpiration in models would be to decipher how vapour pressure deficit and temperature differentially affect the magnitude of daytime and night-time water losses, which has been the topic of other studies^26^. Instead, we here focused on their temporal patterns and observed that neither vapour pressure deficit nor temperature had a significant effect on the pattern of transpiration overnight (Fig. 2). Our results add to an increasing body of research indicating that circadian regulation is important in field settings and acts as a ‘hard clock’ (sensu^22^), meaning that it shows an interactive effect with direct responses to environmental cues (instead of being overridden by them). Circadian regulation is thus no longer a factor we can ignore to understand daily patterns of evapotranspiration.

The substantial water cost of nocturnal transpiration could represent a major problem for agronomic development, especially in areas where water is particularly scarce. It is currently being debated whether or not circadian-induced predawn increases in stomatal conductance ‘prepare’ stomata to respond to daytime environmental cues, ultimately leading to increased carbon uptake and/or stronger stomatal regulation^27,28^. At any rate, circadian regulation of stomatal conductance seems to serve as a mechanism that minimizes the increase in water use overnight, as stomatal conductance is only boosted at the time of minimum atmospheric water demand.

Our study was performed on two species belonging to highly contrasting functional types, and further studies will be necessary to confirm these responses in other species from additional functional groups. However, given the strong influence of circadian regulation, and the widespread occurrence of circadian regulation amongst higher plants^32^, it is to be expected that our results will be applicable to other species as well. It is important to note that even if nocturnal transpiration only accounted for 1% of daytime transpiration globally, it would already have the same impact as documented changes in the intensity of global evapotranspiration with climate change.

## Methods

### Experimental set-up

The experiment was performed in the Macrocosms platform of the CNRS Montpellier European Ecotron. This platform houses 12 identical and independent experimental units. Each unit is composed of a dome under natural light covering a lysimeter inserted in a technical room. The linear series of 12 domes is oriented east-west with two additional domes added at each extremity to eliminate any self-shading edge effects. The 30 m^3^ transparent domes allow for the confinement and control of the atmosphere. Below each dome a lysimeter/ technical room hosts: the soil monolith contained in a lysimeter (2m^2^ area, 2m depth), the lysimeter’s weighing strain gauges and various soil-related sensors, the canopy air temperature and relative humidity conditioning units and the air CO_2_ regulation. Each dome has a circular base area of 25 m^2^, of which 20 m^2^ is covered by concrete and 5 m^2^ central area allocated for the model ecosystems (area 2, 4 or 5 m^2^), height in the centre of the dome is 3.5 m). The airflow from the dome area is prevented to leak into the lysimeter room by the means of fitting metal plates and rubber seals. The concrete is covered with epoxy-resin to prevent its CO_2_ absorption.

Each macrocosm was designed as an open flow gas exchange system. A multiplexer allowed for the CO_2_ concentrations at the inlet and outlet of each dome to be measured every 12 min (LI-7000 CO2/H2O analysers, LI-COR Biosciences, Lincoln, NE, USA). These data combined with the measurement of the air mass flow through each dome allowed for the calculation of canopy carbon assimilation (*A*_c_). Transpiration (mass loss of the lysimeter) was monitored continuously by four CMI-C3 shear beam load cells (Precia-Molen, Privas, France) providing 3 measurements per minute. We ensured only canopy carbon (*A*_c_) and water (*E*_c_) balances were measured by covering the ground with a dark plastic cover that prevented flux mixing. This plastic cover was sealed to the fitting metal plates and not to the lysimeter upper ring. There was a slight over-pressure (+5 Pa) in the dome, and a small proportion of the well mixed air canopy could be passing around the plant stems, therefore flushing the soil respiration and evaporation below the plastic sheet and into the lysimeter room.

The dome was covered by a material highly transparent to light and UV radiation (tetrafluoroethylene film, Dupont USA, 250µm thick, PAR transmission 0.9), and exposed to natural light except during the reduced radiation experiments. Here, an opaque fitted cover (PVC coated polyester sheet Ferrari 502, assembled by IASO, Lleida, Spain) was placed on each dome, and a set of 5 dimmable plasma lamps with a sun-like spectrum (GAN 300 LEP with the Luxim STA 41.02 bulb, Gavita Netherlands), allowed to control radiation. The plasma lamps were then turned off to study dark circadian regulation of stomatal conductance. Our conditions may differ from a cloudy day in that radiation was direct, not diffuse. We were interested in testing how reductions in carbon assimilation affect nocturnal transpiration, therefore, avoiding diffuse radiation was considered advantageous because it increases carbon uptake^33^.

Bean and cotton were planted in rows, one month before the start of the measurements, and thinned at densities of 10.5 and 9 individuals m^−2^ respectively. Six macrocosms were assigned to each species and each individual experiment measuring campaign lasted for 3-4 days. The experiments under constant darkness lasted for 30 hours and we used lysimeter weight readings from three macrocosms per species (six per species in all the other reported experiments). In the three other macrocosms researchers were entering every 4 hours to conduct manual leaf gas exchange measurements at three leaves per dome (LI-6400, LI-COR Biosciences, Lincoln, NE, USA). At the time of measurements, bean and cotton were both at the inflorescence emergence developmental growth stage (codes 51-59 in BBCH scale^34^).

The soil was regularly watered nearly to field capacity by drip irrigation, although irrigation was stopped during the few days of each measuring campaign in order not to interfere with water flux measurements. No significant differences (at *P* < 0.05, paired t-test, n=3) in predawn leaf water potential occurred after a few days of withholding watering. This indicates that no effect of potential changes in soil moisture on plant water status over the course of the experiment.

### Statistical analyses

Transpiration was calculated from the slope of the linear regression between lysimeter weight and time every 3 hours successive periods. Statistical analyses of temporal patterns were then conducted with Generalized Additive Mixed Model (GAMM) fitting with automated smoothness selection^35^ in the R software environment (*mgcv* library in R 3.0.2, The R Foundation for Statistical Computing, Vienna, Austria), including macrocosms as a random factor, and without including outliers (values above 95% quantile during day or night). This approach was chosen because it makes no *a priori* assumption about the functional relationship between variables. We accounted for temporal autocorrelation in the residuals by adding a first-order autoregressive process structure (*nlme* library^36^). Significant temporal variation in the GAMM best-fit line was analysed after computation of the first derivative (the slope, or rate of change) with the finite differences method. We also computed standard errors and a 95% point-wise confidence interval for the first derivative. The trend was subsequently deemed significant when the derivative confidence interval was bounded away from zero at the 95% level (for full details on this method see^37^). Non-significant periods, reflecting lack of local statistically significant trending, are illustrated on the figures by the yellow line portions, and significant differences occur elsewhere.

Differences in the magnitude of total transpiration and in canopy conductance for each species under the different radiation environments were calculated from mixed models that included radiation as a fixed factor (and hour also for canopy conductance) and macrocosms and day of measurement as random factors (each measuring campaign lasted 3-4 days).

## Acknowledgements

This study benefited from the CNRS human and technical resources allocated to the ECOTRONS Research Infrastructures as well as from the state allocation ‘Investissement d’Avenir’ ANR-11-INBS-0001, ExpeER FP7 Transnational Access program, Ramón y Cajal fellowships (RYC-2012-10970 to VRD and RYC-2008-02050 to JPF) and internal grants from UWS-HIE to VRD and ZALF to AG. We remain indebted to E. Gerardeau, D. Dessauw, J. Jean, P. Prudent (Aїda CIRAD), C. Pernot (Eco&Sol INRA), B. Buatois, A. Rocheteau (CEFE CNRS), S. Devidal, C. Piel, O. Ravel and the full Ecotron team, J. del Castillo, P. Martίn, A. Mokhtar, A. Pra, S. Salekin, J. Voltas (UdL), S. Garcίa-Muñoz (IMIDRA), Z. Kayler and K. Pirhofer-Walzl (ZALF) for outstanding technical assistance during experiment set-up, plant cultivation or subsequent measurements.

### Author Contributions

V.R.D., J.R and A.G. designed research with input from J.P.F. and A.M. V.R.D. and J.G.A. analyzed the data. V.R.D. wrote the first draft. All authors performed research and contributed to revisions.

### Additional Information

#### Competing financial interests

The authors declare no competing financial interests.

